# qSR: A software for quantitative analysis of single molecule and super-resolution data

**DOI:** 10.1101/146241

**Authors:** J. Owen Andrews, Arjun Narayanan, Jan-Hendrik Spille, Won-Ki Cho, Jesse D. Thaler, Ibrahim I. Cissé

## Abstract

Single-molecule based super-resolution microscopy (Betzig et al., 2006; Hess et al., 2006; Rust et al., 2006) provides insight into the spatiotemporal organization and dynamics of biomolecules at the length scale of ten to hundreds of nanometers, which is most relevant to protein clustering in a broad range of cellular processes. A series of quantitative analysis approaches have been developed to infer the characteristic size of protein clustering via pair correlation analysis (Sengupta et al., 2013; Sengupta et al., 2011; Veatch et al., 2012) or to merge multiple blinking localizations from single fluorophores (Annibale et al., 2011), to identify clusters based on density of detections (Ester, 1996), and to measure temporal dynamics of protein clustering in living cells (Cho et al., 2016a; Cisse et al., 2013). At the same time, laboratories are increasingly gaining access to the instrumentation of super-resolution imaging by means of commercial microscopes or dedicated imaging centers on research campuses. To concomitantly increase access to the analysis algorithms, we have developed qSR, an integrated software package for super-resolution data analysis.

In qSR, we have implemented established algorithms for pair-correlation analysis and spatial clustering, including a new application of FastJet (Cacciari et al., 2012; Dokshitzer et al., 1997; Wobisch and Wengler, 1999), a cluster analysis package developed by the particle physics community. A major feature in qSR, which to our knowledge has not been present in previous super-resolution analysis packages (Andronov et al., 2016; Levet et al., 2015; Malkusch and Heilemann, 2016; Pengo et al., 2015), is the integrated toolset to analyze the temporal dynamics underlying live cell super-resolution data.

The pointillist data obtained from single-molecule based super-resolution microscopy techniques—such as photoactivated localization microscopy (PALM) (Betzig et al., 2006; Hess et al., 2006), stochastic optical reconstruction microscopy (STORM) (Rust et al., 2006) and direct STORM (Heilemann et al., 2008)—can be imported into qSR for visualization and analysis (**Figure 1B**). Super-resolution images can be reconstructed, and represented in a red-hot color coded image, by convolving the point pattern of detections with a Gaussian intensity kernel corresponding to the localization uncertainty (**Figure 1C**).

**Figure 1:**
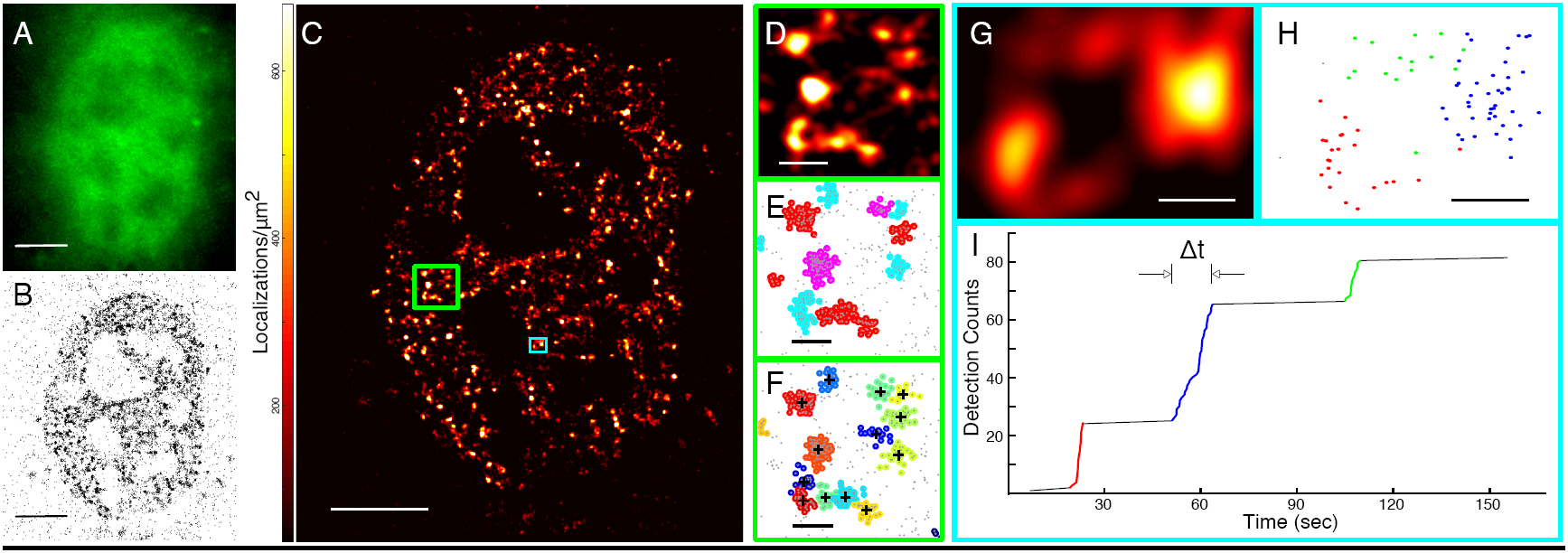
qSR facilitates analysis of the spatial organization and temporal dynamics of proteins in live cell super-resolution data. (**A**) Conventional fluorescence image of a nuclear protein inside a living cell. Data is cropped to an area around the cell nucleus where the protein (in this example RNA Polymerase II) is primarily localized. (**B**) A pointillist image (point representation of the set of detections) of single-molecule localizations is represented. Singe-molecule localization data is the input for the qSR software. (**C**) A super-resolution reconstruction image is represented using qSR. In this figure, each localization is represented as a Gaussian intensity distribution with a standard deviation of 50 nm. A red-hot color code is used to represent the local density detections. (**D** & **E**) Spatial clustering of the data within the region highlighted in the green box shown in (**C**) is performed using the DBSCAN algorithm embedded in qSR. DBSCAN allows for the detection of clusters of varying shapes and sizes, while excluding noise below a given density threshold. (**F**) Spatial clustering of the same region—in the green box shown in (**C**)—is performed using the FastJet algorithm embedded in qSR. (**G**, **H** & **I**) Time-correlation super-resolution analysis (tcPALM) reveals temporal dynamics within any region of interest (ROI)—here ROI shown in (**G**), and highlighted in cyan in (**C**). In (**I**), for the selected ROI, a plot of the cumulative number of localizations as a function of time is represented. Localizations are not randomly distributed in time but come in bursts. Three temporal clusters are shown in red, blue and green. As illustrated for the example highlighted in blue, the apparent cluster lifetime (Δt) is the span of the clustering event in the time axis, and the burst size is the span in the detection counts axis. Localizations belonging to the three temporal clusters highlighted in (**I**) are plotted spatially in their corresponding (red, blue, green) colors in (**H**). Clusters of localizations which are grouped by time in (**I**) are also distinctly clustered in space. Scale Bars: A, B, & C – 5 μm; D, E & F – 500 nm G & H – 200 nm.

In addition, qSR enables the quantitative analysis of the spatial distribution of localizations. The qSR analysis tools provide the user with both a summary of detected clusters, including their areas and number of detections, and a global metric of the distribution of sizes via the pair correlation function. For identifying spatial clusters, we have implemented both centroid-linkage hierarchical clustering using FastJet (Cacciari et al., 2012; Dokshitzer et al., 1997; Wobisch and Wengler, 1999) illustrated in **Figure 1E**, and density-based spatial clustering of applications with noise (DBSCAN) (Ester, 1996) as illustrated in **Figure 1D**.

In super-resolution data, thousands of localizations are typically recorded for an image, leading to long analysis time. In developing FastJet, Cacciari and co-workers exploited advances in computational geometry and data structures to develop a version of hierarchical clustering with an asymptotic runtime speedup compared with previous implementations of hierarchical clustering. This version of hierarchical clustering is sufficiently fast to use at the Large Hadron Collider for on-line reconstruction of data collected at megahertz rates (Cacciari et al., 2012). Via the qSR software, FastJet can analyze a typical super-resolution dataset within a few seconds (**Supplemental Information Section B.2.b**). By storing the full tree structure, the user can quickly re-cluster data and compare the resulting clusters at varying characteristic sizes. DBSCAN (Ester, 1996), on the other hand, was developed to find clusters in noisy data. The local density at each point is calculated as the number of adjacent points within a local neighborhood. Points below a density threshold are designated as noise, and the remaining points are joined together into clusters. The clustering algorithms for FastJet and DBSCAN implemented in qSR are housed within an easy to use graphical interface, and gives the user immediate visual feedback on the quality of clustering.

qSR also integrates a number of tools to facilitate the discrimination between single molecule oversampling and true protein organization. An individual fluorophore may be visible for multiple, intermittent frames, leading to multiple localizations of single molecules. Such an oversampling of single molecules could appear as artefactual spatial clusters. One tool, pair-correlation PALM (pcPALM), developed by Sengupta *et al*. (Sengupta et al., 2011) addresses this problem. pcPALM is a test that can decompose the spatial correlation into the contribution from single molecule photo-physics and the contribution from the protein clusters by using the pair correlation function. qSR provides a means for simple calculation of the pair correlation function (**Supplemental Information Section B.2.a**). While pcPALM was initially implemented for inferring spatial clustering of membrane proteins, we find that it can be readily expanded for spatial clusters within the cell body including in the cell nucleus (Cho et al., 2016a; Cisse et al., 2013). Moreover, to reduce the number of localizations from a single molecule, Annibale and coworkers propose merging localizations that occur within a user-defined spatial and temporal threshold separation (Annibale et al., 2011). In qSR, this is implemented as a “spatiotemporal merging” filter. pcPALM and spatiotemporal merging can be readily performed together in qSR to facilitate the positive identification of protein clusters.

Finally, qSR adopts time-correlated super-resolution analyses—for example tcPALM (Cho et al., 2016a; Cisse et al., 2013)—to measure the dynamics of sub-diffractive protein clustering in living cells. In live cell super-resolution data, when clusters assemble and disassemble dynamically, the plots of the temporal history of localizations in a cluster show temporal bursts of localizations (**Figure 1G-I**). The apparent cluster lifetime and burst size can then be measured, and other clustering parameters, including clustering frequency, can be calculated (Cho et al., 2016a; Cisse et al., 2013). For a sample data set, and step by step instruction on how to perform tcPALM please see **Supplemental Information Section B.1**. It is important to ensure that apparent bursts of detections are not due to long-lived single molecules. Therefore, at minimum, control experiments with fixed cells expressing the fluorophore alone (i.e. unfused to any other protein) should be performed to characterize the temporal profile of individual molecules (**Supplemental Information Section B.3**).

In summary qSR integrates complementary algorithms that together form a unique tool for the quantitative analysis of single molecule based super-resolution —PALM (Betzig et al., 2006; Hess et al., 2006) and STORM (Rust et al., 2006)— data from both fixed and living cells. The input for qSR is a single-molecule localization dataset, and the prior image processing can be performed with popular open-source softwares like ImageJ (Abràmoff et al., 2004; Collins, 2007; Schneider et al., 2012). qSR readily accepts as inputs the files generated by super-resolution localization plug-ins in ImageJ, including QuickPALM (Henriques et al., 2010), or Thunderstorm (Ovesný et al., 2014) which are freely available as add-ons to ImageJ. We note that there are other super-resolution approaches, such as STED (Hell and Wichmann, 1994; Klar et al., 2000), and structured illumination microscopy (Gustafsson, 2000, 2005) that are not fundamentally based on single-molecule localizations, and the current implementation of qSR is not adapted for those datasets.

qSR is provided as an open source software package with a graphical user interface that a researcher can download and use directly. The live cell super-resolution data represented in **Figure 1** is provided as sample data (Cho et al., 2016b) packaged along with the software. Installation instructions and protocols for performing the analyses are provided as Supplemental Information. qSR is maintained and distributed for free on www.github.com/cisselab/qSR. By providing qSR as an open source platform, we hope to encourage other users in the community to develop and integrate any multitude of algorithms, features and modules for qSR, as the field evolves. Third party algorithms can be readily linked via the qSR portal on Github, and users are prompted to directly cite the original publications for any third party algorithm they use within the qSR platform.

## Acknowledgments

We thank Will Conway (MIT) and Namrata Jayanth (MIT) for helpful comments. JOA is supported by the National Science Foundation Graduate Research Fellowship. Any opinion, findings, and conclusions or recommendations expressed in this material are those of the author and do not necessarily reflect the views of the National Science Foundation. The work is also supported by the National Institutes of Health, the National Cancer Institutes through the NIH Director’s New Innovator Award, DP2-CA195769 to IIC and funds from MIT Department of Physics. Further support for this work was provided by NIH 4D Nucleome program, grant U54- DK107980. The content is solely the responsibility of the authors and does not necessarily represent the official views of the National Institutes of Health. The software is distributed for free and open source under the GNU General Public License (GPLv2 / LGPLv2) Version 2.

